# Do the shuffle: Expanding the synthetic biology toolkit for shufflon-like recombination systems

**DOI:** 10.1101/2024.10.25.620183

**Authors:** Jan Katalinić, Morgan Richards, Alex Auyang, James H. Millett, Manjunatha Kogenaru, Nikolai Windbichler

## Abstract

Naturally occurring DNA inversion systems play an important role in the generation of genetic variation and adaptation in prokaryotes. Shufflon invertase (SI) *Rci* from plasmid R64, recognizing asymmetric *sfx* sites, has been adopted as a tool for synthetic biology. However, the availability of a single enzyme with moderate rates of recombination has hampered the more widespread use of SIs. We identified 14 previously untested SI genes and their *sfx* sites in public databases. We established an assay based on single-molecule sequencing that allows the quantification of the inversion rates of these enzymes and determined cross-recognition to identify orthogonal SI/*sfx* pairs. We describe SI enzymes with substantially improved shuffling rates when expressed in an inducible manner in *E. coli*. Our findings will facilitate the use of SIs in engineering biology where synthetic shufflons enable the generation of millions of sequence variants *in vivo* for applications such as barcoding or experimental selection.

## Introduction

Bacteria employ multiple mechanisms for DNA inversion to achieve phenotypic heterogeneity within the population^1^, a process that is now believed to be more widespread than previously thought^2^. This strategy generates variation and allows the selection of lineages with a set of beneficial genetic traits, for instance under conditions of environmental stress. One mechanism to achieve this is conservative site-specific DNA recombination mediated by a recombinase which typically binds to a single pair of recognition sites and inverts the intervening DNA sequence. However, a more interesting case is presented by shufflons - multiple inversion systems that contain several recombination sites that flank and separate multiple invertible coding regions and as a result can produce a larger set of alternative proteins^3^. The presence of multiple recombination sites enables a single recombinase or shufflon invertase (SI) to ‘shuffle’ between DNA segments to produce alternative alleles. In *Bacteroides fragilis* the SI Tsr0667 shuffles a gene cluster of outer-membrane proteins of the SusC/SusD family to produce alternative alleles utilising different promoters^4^. This mechanism leads to varied combinations and expression levels of SusC/SusD proteins contributing to differential polysaccharide utilisation of *B. fragilis*. Other multiple inversion systems include those responsible for antigen variation by alternating expression of *vsa* gene variants in *Mycoplasma pulmonis*^5^, shuffling outer membrane protein genes in *Dichelobacter nodosus*^3^ or immune evasion in *Campylobacter fetus* by varying surface layer protein genes^6^.

One of the best characterised systems is the shufflon of the pil operon on plasmid R64^7^, a DNA inversion system that functions as a biological switch to select between alternative C-terminal segments of the pilV protein^8,9^. Conjugative thin pili which are required for liquid matings and the *pilV* products are tip-located adhesins of the type IV pilus that recognize lipopolysaccharides of recipient bacterial cells and thus determine recipient specificity in matings. The pil operon harbours the *rci* gene, coding for shufflon invertase *Rci*, and the adjacent shufflon made of four DNA segments separated and flanked by seven recognition sites, also known as shufflon crossover or *sfx* sites, specifically recognized by *Rci*^10,11^. *Rci’s sfx* sites are composed of a 12bp left arm, a 7bp core or spacer sequence, and a 12bp right arm. Fully conserved nucleotides are found only in the core and right arm whereas the sequence of the left arm is highly variable ^10^. It has been proposed that this asymmetry predetermines *Rci* binding and limits its activity to perform inversions only. The model suggests that *Rci* bound to the right arm recruits a second *Rci* monomer to bind to the left arm sequence non-specifically and is supported by experiments with artificially symmetric *sfx* sites that yield both inversions and excisions events. This cooperative binding is thought to be mediated by an additional C-terminal domain not present in related Tyrosine recombinases like Cre^12^ which perform both excisions and inversions.

It has been demonstrated that *Rci*-mediated shuffling in *E. coli* requires no co-factors making it an attractive synthetic biology tool for sequence diversification in different contexts^8^. A proof-of-principle study for the use of DNA inversion systems to enable *in vivo* genetic barcoding combined *Rci* with artificial 5-module and 11-module shufflons yielding a theoretical maximum of 384 and 176,947,200 barcodes respectively^13^. The authors used long-read sequencing to explore the unique barcodes generated by this system and demonstrated *Rci*’s potential for generating a large amount of sequence diversity within a synthetic construct. Whilst the literature thus suggests that *Rci* is capable of shuffling synthetic constructs in *E. coli*, its efficiency in doing so is inconsistent and requires long incubation periods^8,13^. *Rci* has also been demonstrated to show low activity in eukaryotes^14,15^, including the occurrence of deletions not previously observed in studies on wild-type *Rci*. Recently an approach that employed directed evolution managed to generate *Rci* mutants with a higher inversion frequency and a negligible rate of deletion in eukaryotes including human HEK293 cells^15^. The study did not test how these *Rci* mutants behave in the bacterial context (Y. Wu, personal communication).

Together this suggests that the availability of a greater range of SIs and variant *sfx* sites could provide more enzymes suitable for particular applications, contexts, and organisms and additional starting points for evolving enzymes for custom synthetic biology applications. It could also provide the starting point for the establishment of orthogonal shuffling systems or yield SI enzymes with higher rates of activity especially in the context of larger synthetic shufflons. We thus sought to identify variant SI enzymes homologous to *Rci* recognizing differing *sfx* sites and to establish a reporter system that allowed the comparison of their shuffling capabilities.

## Results and Discussion

To identify a set of novel SI enzymes we relied on the fact that in natural DNA inversion systems SI genes and shufflons are typically co-located in close proximity. To that end, we took an iterative DNA sequence mining approach (Figure 1A) starting with the amino acid sequence of *Rci* (WP_001139955) in a query against predicted DNA translations of sequences in the NCBI core nucleotide database ‘core_nt’. We extracted putative SI protein sequences as well as up to 5kb of the DNA regions flanking each genomic or plasmid hit. Flanking DNA sequences were then screened for 13-mers which occurred multiple times or occurred at least twice as inverted repeats. This is based on the longest conserved subsequence within *Rci’s* 31-nt *sfx* site of 13 nucleotides^10^. Organisms with hits lacking identifiable repeats or hits with repeats identical to the conserved portion of the original *sfx* site, were excluded from the database in the subsequent iterative searches. The protein sequences of SIs featuring *sfx* repeat sequences non-identical to *Rci*’s *sfx* site were then used as seeds to initiate additional search iterations. In total, out of 314 DNA loci which shared translated sequence homology with *Rci*, 94 contained at least two 13-nt direct repeats, of which 56 also had at least one inverted repeat. Of those, 14 were non-identical to *Rci’s sfx* site. Two homologs with palindromic repeats were not considered due to the lack of asymmetry in their cognate recognition sites. For simplicity, here we named *Rci* homologs according to their species of origin in the following format: SI-species name. Of the 14 homologs, one was discarded since its protein sequence was significantly shorter than the other SIs, and another because its sequence was identical to *SI-E. E76*. In addition to these 12, two further homologs had already been identified in a prior database screen (*SI-H. paralvei* and *SI-Y. pseudotuberculosis*). We confirmed using AlphaFold modelling that all 14 considered homologs have the additional, flexible C-terminal domain like Rci which is not present in Cre (Figure 1B) or other Tyrosine recombinases such as XerC, XerD, HP1, and λ^16^. We also confirmed the presence of the catalytic tetrad Arg-His-Arg-Tyr^16,17^ in these homologs (Supplementary Figure 1). Furthermore, we included three *Rci* mutants from Han, P. et al. (2021) to be tested. Mutant m8 was reported to have the highest inversion to deletion ratio in *Saccharomyces cerevisiae*, while mutants m18 and m34 were reported to have the highest inversion rates in the same organism^15^. Thus, a total of 17 SIs were evaluated alongside the original *Rci* in our experimental system (Table 1, Supplementary File 1). A single *sfx* site was defined for each enzyme picking, where possible, the most common sequence variant for the variable left arm or by randomly choosing the variable arm from a single *sfx* repeat.

**Table 1:**
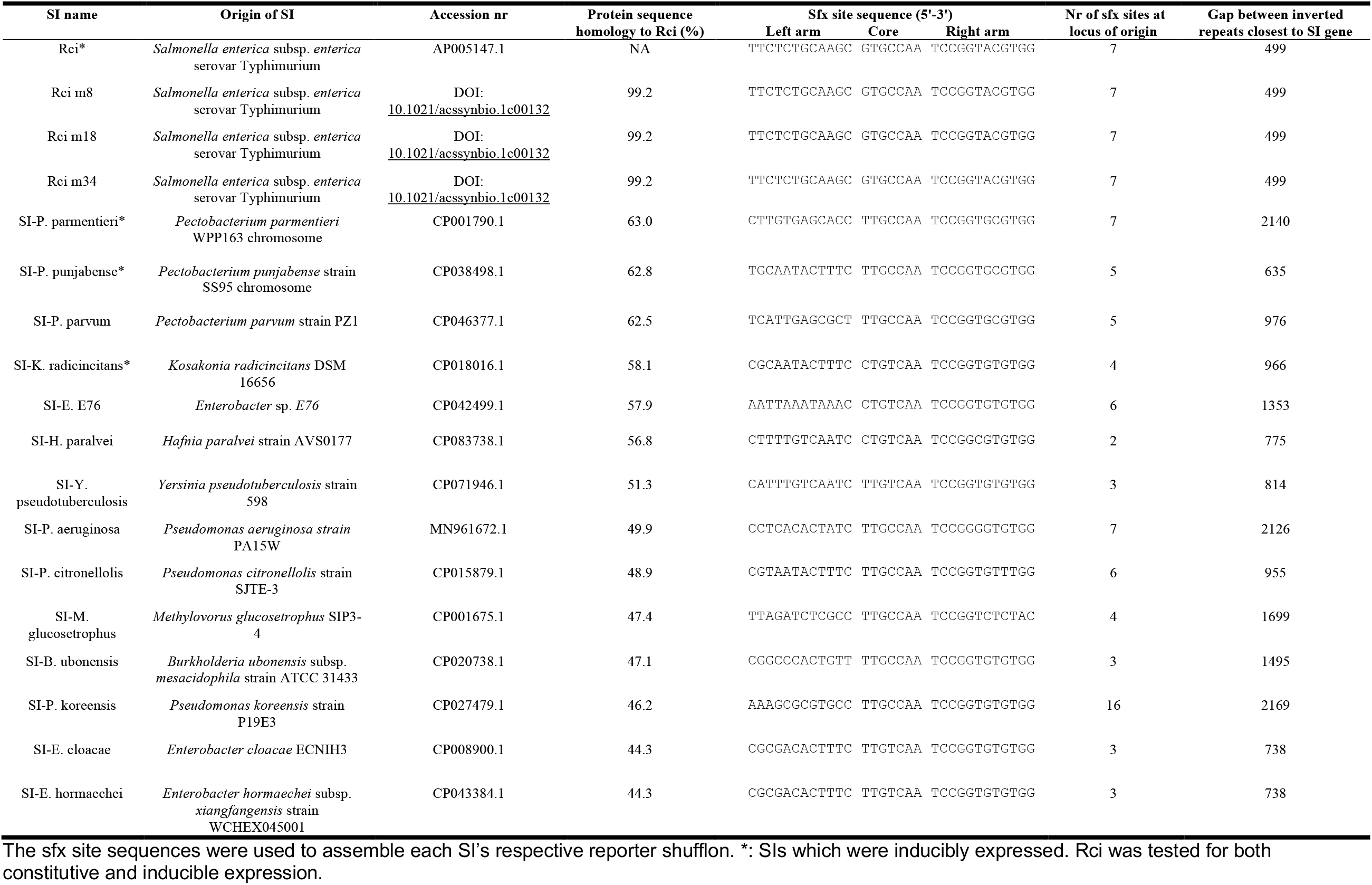
Tested SIs and their cognate sfx site sequences.

**Figure 1.**
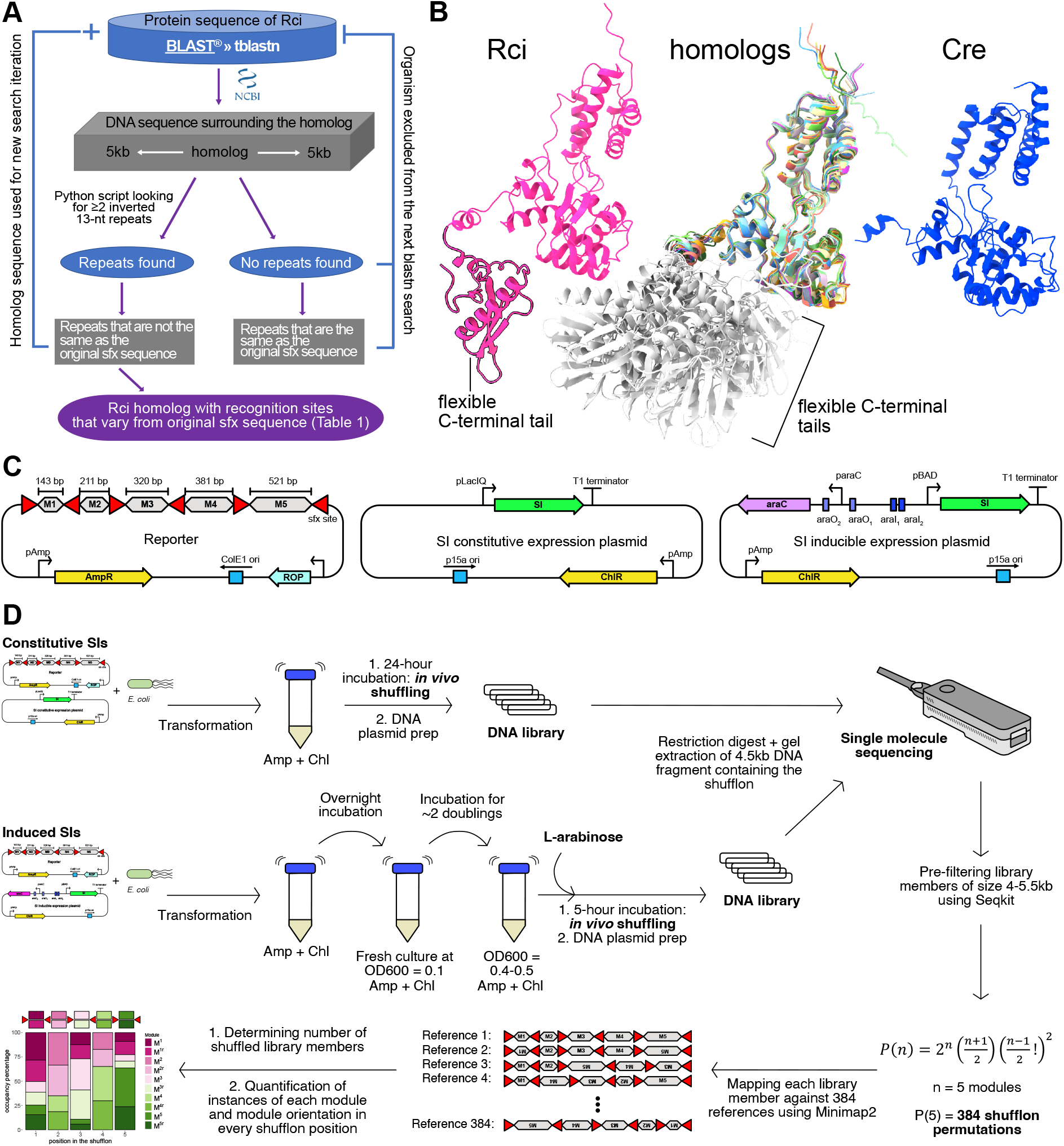
Experimental overview. **A**. Bioinformatic strategy for the isolation of shufflon invertases (SIs) and their recognition sites based on homology to *Rci*. **B**. Comparison of AlphaFold protein models of *Rci*, superimposed homologs, and *Cre*. The additional C-terminal domain is outlined in black for *Rci*, and displayed in light grey for homologs. **C**. Structure of the reporter (left) as well as the constitutive (middle) and inducible (right) expression plasmids used in this study. AmpR: ampicillin resistance gene. ROP: repressor of primer. ChlR: chloramphenicol resistance gene. M1 to M5 represent the invertible dummy sequences. **D**. Experimental workflow for both inducible and constitutively expressed SIs.

Next, we established a standardised system for comparing Rci’s shuffling capabilities with the other 17 SIs. It has been reported that in *E. coli* expressing *Rci* from low copy number plasmids prevents protein aggregation^13^. Our initial attempts also showed that the *rci* gene or its promoter acquire mutations when *Rci* is overexpressed from high-copy vectors suggesting some level of toxicity (data not shown). All synthesised SI’s open reading frames were therefore cloned into a low copy vector (Figure 1C) with the p15a origin of replication under the transcriptional control of the pLacIQ promoter (Registry of Standard Biological Parts: BBa_K1695000). However, out of the 18 assembled SIs three (*SI-P. punjabense, SI-P. parmentieri*, and *SI-K. radicincitans*) were found to accumulate mutations even in the low-copy expression vector as evident during standard passaging and plasmid sequencing. They were therefore cloned into an L-arabinose-inducible^18^ expression vector (Figure 1C) alongside *Rci*.

Next, we assembled, for each *sfx* variant a reporter plasmid harbouring a five-module shufflon of “dummy” sequences of increasing length (143 to 521bp) flanked by six recombination sites in alternating directions (Figure 1C). Since shuffling of 100 bp fragments with *Rci* had been successfully demonstrated before^13^, we chose our minimum length to be close to that value. Each module can be inverted in place, and, following inversion, is expected to be able to occupy any other position of the same parity: even modules can occupy any even position, odd modules can only occupy any odd positions for a total of 384 possible shufflon permutations.

Figure 1D illustrates the experimental workflow for our shuffling assay. First, each SI was co-transformed with its cognate reporter into Stbl3 *E. coli* cells followed by liquid culture in the presence of ampicillin and chloramphenicol to select for both reporter and SI expression plasmids. For constitutively expressed SIs, the cultures were incubated for 24 hours (Figure 1D) following Peikon et al. who demonstrated *Rci* shuffling of multi-modular cassettes after overnight incubation of their *E. coli* liquid culture^13^. For inducibly expressed SIs, cultures were first incubated overnight and then used to inoculate fresh cultures at an optical density OD600 of 0.1 units. After roughly two doublings, L-arabinose was added for a final concentration of 0.2% upon which the cultures were incubated for a further 5 hours. The plasmid DNA library was then extracted from each culture and subjected to a restriction digest to isolate the shufflon fragments. The DNA shufflon libraries were then barcoded for pooled single nucleotide sequencing using the Oxford Nanopore Technologies (ONT) MinION system. Following basecalling and deconvolution, each library was mapped against the 384 possible reference permutations that can be generated by each reporter shufflon of five modules. For further details describing the bioinformatic analysis, see Materials & Methods.

Figure 2A shows the mean shuffling rates, i.e. the fraction of shufflon copies not in the original (i.e. starting) configuration over two biological replicates for each constitutively-expressed SI tested. Five SIs produced at least one replicate with a shuffling rate above 50%: wild-type *Rci*, homologs *SI-E. E76* and *SI-B*.*ubonensis*, and *Rci* mutants m18 and m34. Of those, wild-type *Rci* displayed the highest mean shuffling rate of 97% followed by *SI-E. E76* and *SI-B. ubonensis* with 79% and 74% mean shuffling rates respectively. All other homologs exhibited either no shuffling or a minimal amount of up to about 7%. Of the three *Rci* mutants, mutant m8 yielded no shuffling whilst mutant m18 yielded the highest mean shuffling rate. Figure 2B shows the module occupancy after shuffling for those SIs with an appreciable shuffling rate as well as *SI-P. citronellolis* as an example of an unshuffled library. We observed a previously reported bias, namely that terminal modules are less frequently shuffled than central modules flanked by multiple recombination sites. The shuffling rates of homologs *SI-E. E76* and *SI-B. ubonensis* did not exhibit statistically significant differences in pairwise comparisons to wild-type *Rci* (Dunnett’s C test). However, a closer look at the obtained reporter libraries shows that these SI’s also generated fewer library members with more than 1 inversion (Supplementary Figure 2A) and the libraries covered a smaller share of the total diversity possible (384) in an analysis of 4000 randomly selected library members per SI (Supplementary Figure 2B). Interestingly, even though *SI-E. E76* yielded a higher mean shuffling rate than *SI-B. ubonensis*, the *SI-B. ubonensis* library displays more evenly distributed module occupancies in every position (Figure 2B). It also yielded 259 out of 384 permutations of its reporter shufflon while *SI-E. E76* yielded only 217 permutations (Supplementary Figure 2B). Similarly, despite *Rci* m18’s lower mean shuffling rate compared *SI-E. E76* and *SI-B. ubonensis*, and obvious biases in module occupancy (Figure 2B), it generated 293 different permutations. These findings suggest that a high shuffling rate may not always reflect a library’s diversity. The wild-type *Rci* library showed the most evenly distributed configuration of shufflon positions (Figure 2B). Among the 4000 randomly sampled library members from both biological replicates, 373 out of 384 permutations were identified (Supplementary Figure 2B). Therefore, among the constitutively expressed SIs, it appears to be the most active SI and generated the most diverse library.

**Figure 2.**
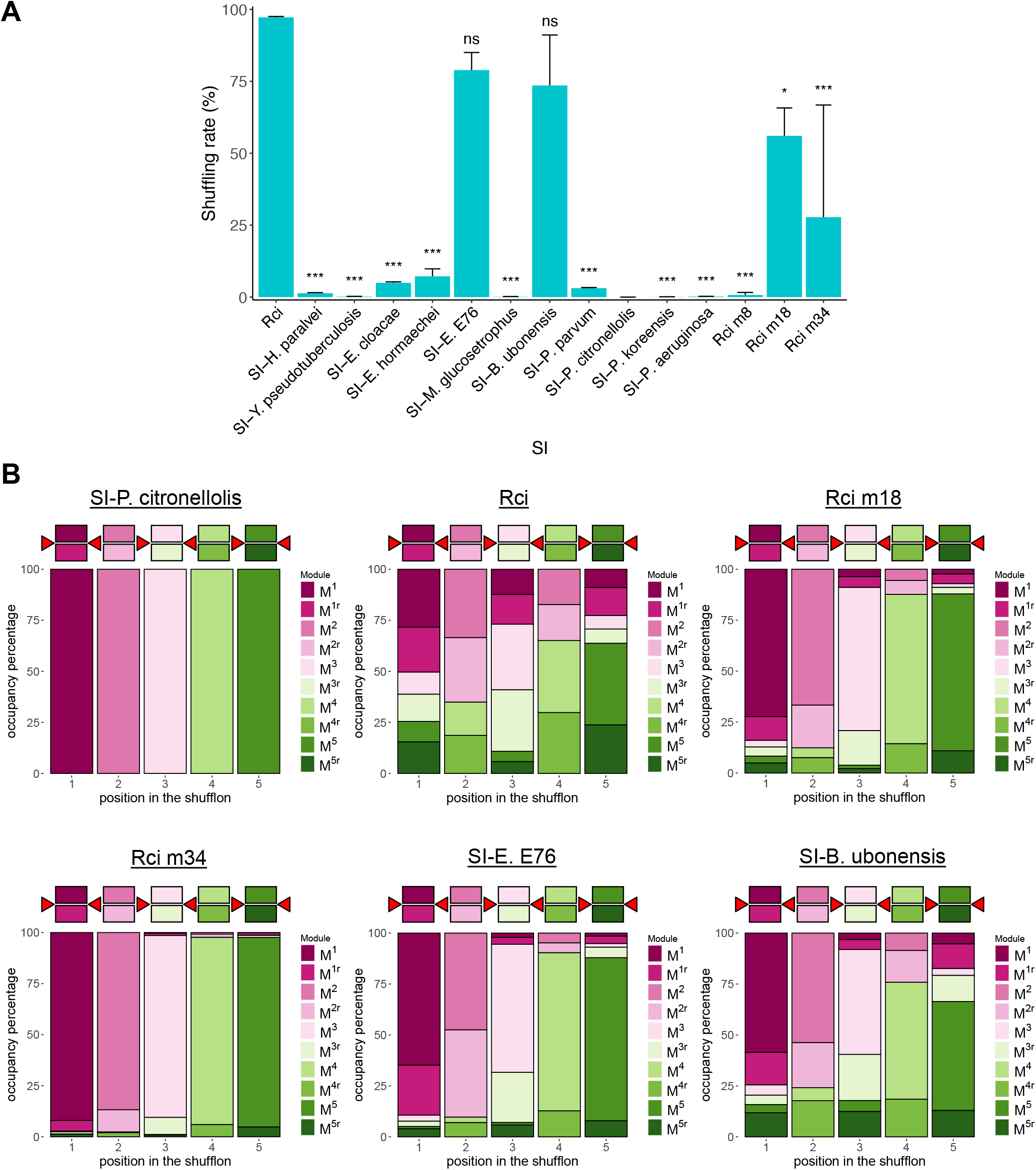
Shuffling of reporters by constitutively expressed SIs. **A**. Bar graphs showing the mean shuffling rate between two biological replicates. Each estimate is based on a minimum of 1500 analysed reads. Error bars represent standard deviation (SD). Following ANOVA, Dunnett’s C test was performed for pairwise comparisons to *Rci*. p-values are represented by *, 0.05 ≥ p > 0.01; **, 0.01 ≥ p > 0.001; ***, 0.001 ≥ p; ‘ns’, non-significant. **B**. Indicative module occupancy in the shuffled library of selected SIs. The diagram above each plot represents the shufflon in its reference (unshuffled) form. Coloured squares represent the dummy sequences when present on sense or antisense strands and red triangles indicate the *sfx* sites. For each panel, 6000 library members were randomly sampled and pooled from two biological replicates.

Next, we aimed to determine whether orthogonality between SIs existed using our set of active enzymes and reporters with different *sfx* variants. An orthogonal system of SIs would be a useful addition to the toolkit, as it could provide precise spatial and temporal control of SI-mediated inversions or could be used to establish dual index barcodes shuffled by different SIs at different times. For this purpose *Rci*, and the two most active homologs, *SI-E. E76* and *SI-B*.*ubonensis*, were tested for shuffling each other’s reporters, as well as for shuffling their cognate reporters as controls. Additionally, they were tested for shuffling reporters with *sfx* sites most dissimilar to their own. The result of this experiment is shown in Figure 3. Overall, shuffling rates were lower for all 3 SI’s tested in this experiment with *SI-B. ubonensis* showing the most substantial decrease of shuffling its own reporter (by 50% on average). *SI-B. ubonensis* was the only one out of the three SIs that did not shuffle any of the non-cognate reporters at an appreciable rate (Figure 3A). Surprisingly, *SI-E. E76* exhibited high shuffling activity with 3 different non-cognate reporters, in all three cases surpassing the rate for its cognate reporter. There is no statistical difference between mean shuffling rates of *SI-E. E76* with *E. cloacae* and *B. ubonensis* reporters compared to *Rci*’s shuffling rate for its cognate reporter (two-sample t-tests with Welch’s correction) which is the combination that had previously yielded the most diverse reporter library. This is supported by the module occupancy graphs across the reporter shufflons for both *E. cloacae* and *B. ubonensis* (Figure 3B). Whilst there is a statistically significant difference between the mean shuffling rates of *Rci* and *SI-E. E76* for various reporters, our analysis did not yield a fully orthogonal pair of SIs in combination with reporters harbouring different *sfx* variants.

**Figure 3.**
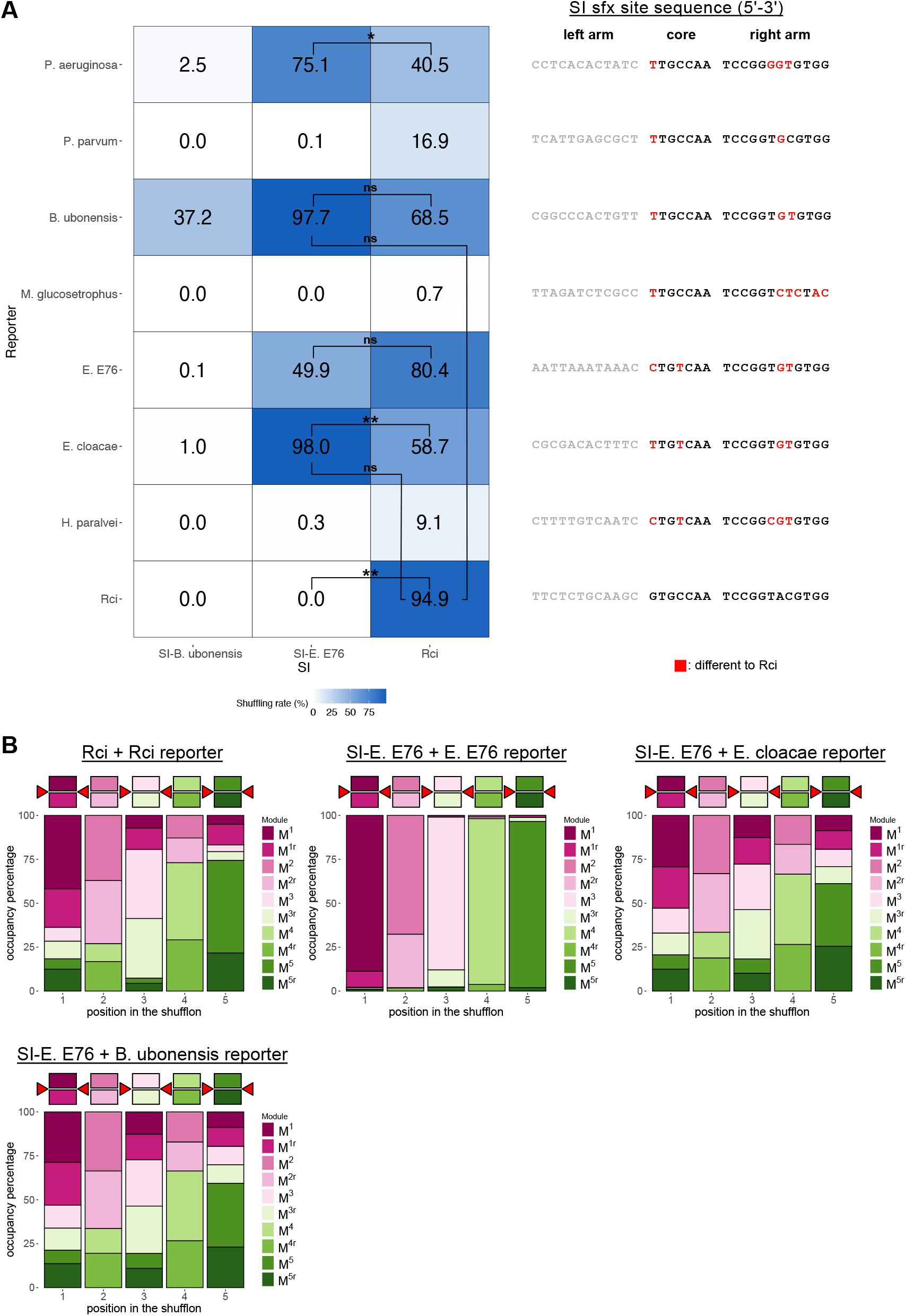
Testing constitutively expressed SIs for orthogonality and cross-recognition. **A**. Each panel displays the mean shuffling rate from two biological replicates for each corresponding SI + reporter pair. An average of 6093 reads were analysed per panel. Two-sample t-tests with Welch’s correction were performed for pairwise comparisons of mean shuffling rates. p-values are represented by *, 0.05 ≥ p > 0.01; **, 0.01 ≥ p > 0.001; ***, 0.001 ≥ p > 0.0001; ****, p ≤ 0.0001; ‘ns’, non-significant. **B**. Indicative module occupancy in the shuffled library of selected SIs. The diagram above each plot represents the shufflon in its reference (unshuffled) form. Coloured squares represent the dummy sequences when present on sense or antisense strands and red triangles indicate the *sfx* sites. For each panel, 6000 library members were randomly sampled and pooled from two biological replicates.

The results of the experiments with inducibly expressed SIs are summarised in Figure 4. In accordance with expectations, *Rci* induced no significant shuffling of its reporter in the absence of L-arabinose in the culture. Following induction with L-arabinose, the mean shuffling rate observed after 5 hours was 42.1% for *Rci*. The other 3 SI variants also generated shuffling of their cognate reporters. 2 of them, *SI-P. punjabense* and *SI-K. radicincitans*, produced mean shuffling rates of 89.8% and 93.2% respectively for their cognate reporters, significantly outperforming *Rci* under these conditions (Dunnett’s C test for pairwise comparisons, Figure 4A). *SI-K. radincincitans* in particular displayed a more balanced module occupancy (Figure 4B) and generated the most diverse library (370 out of 384 possible shufflon permutations) within the 4000 randomly sampled library members (Supplementary Figure 2B) compared to *Rci* (95 out of 384). It has previously been demonstrated that the 563 bp long segment A of the original R64 shufflon had the highest inversion frequency. Shortening it by about 100 bp and 250 bp had no significant impact on the inversion frequency, while increasing the sequence length by approximately 2.5 times as well as 3.5 times reduced its inversion frequency^8^. We therefore investigated the impact of module length on SI inversion frequency in our experiments. We considered the data of all five constitutively expressed SIs that we found to be active as well as all inducibly expressed SIs. From each library we exclusively considered reads which represented the original reference order of modules and quantified the inverted and uninverted configurations of each module. The reason for this selection was to consider library members which were likely the products of only a single module inversion. A multiple regression analysis was carried out to assess the effect of SI and module length on the module inversion frequency across SI’s (Supplementary Figure 3). It suggests a decline in inversion frequency with increasing module length (module length being a significant coefficient p<0.00361) and confirms the expected significant effect of the SI employed (p<1.484×10^−9^) on inversion efficiency.

**Figure 4.**
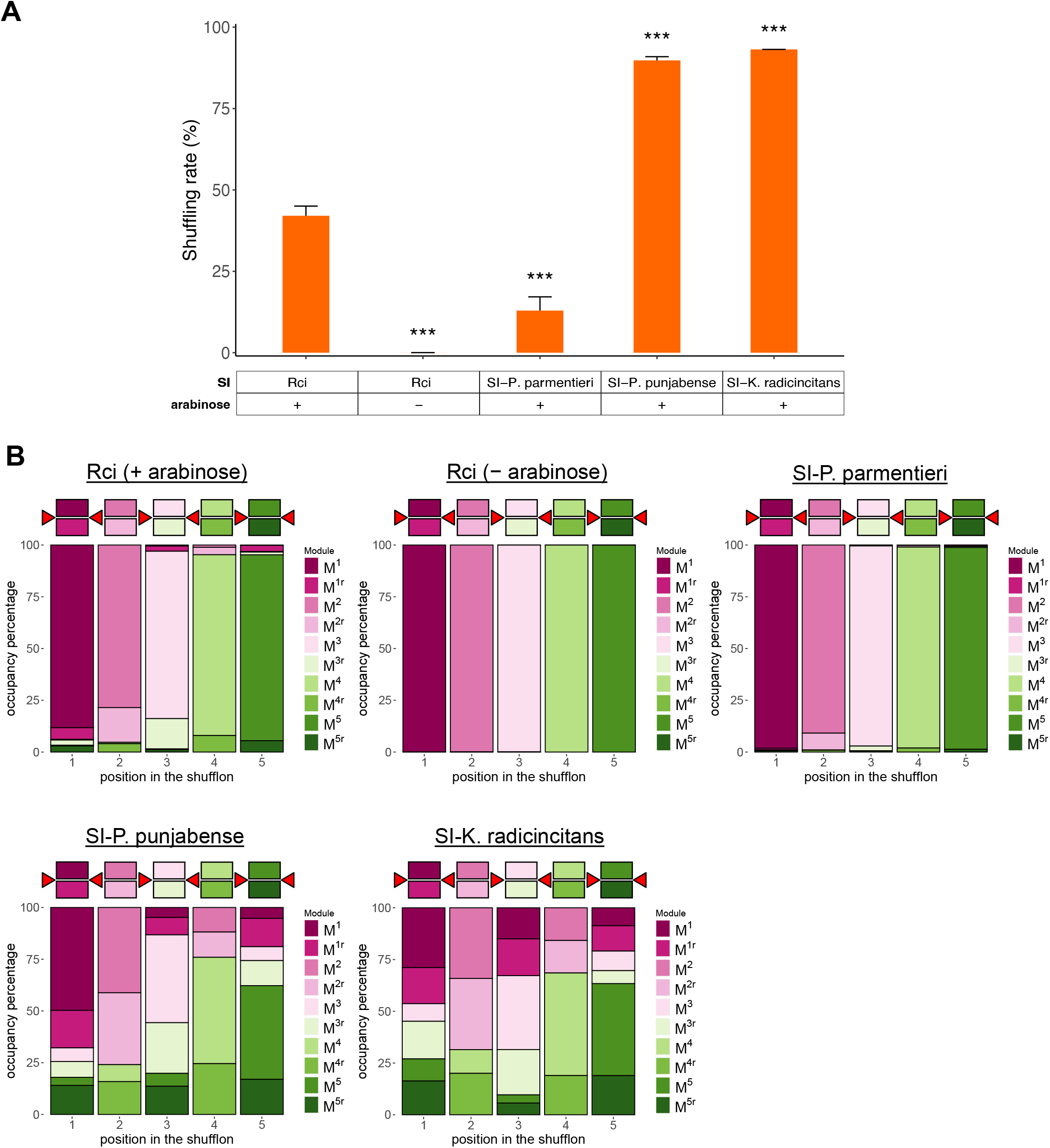
Shuffling of reporters by inducibly expressed SIs. **A**. Bar graphs showing the mean shuffling rate between two biological replicates. Each estimate is based on a minimum of 1300 analysed reads. Error bars represent standard deviation (SD). Following ANOVA, Dunnett’s C test was performed for pairwise comparisons to *Rci*. p-values are represented by *, 0.05 ≥ p > 0.01; **, 0.01 ≥ p > 0.001; ***, 0.001 ≥ p; ‘ns’, non-significant. **B**. Indicative module occupancy in the shuffled library of selected SIs. The diagram above each plot represents the shufflon in its reference (unshuffled) form. Coloured squares represent the dummy sequences when present on sense or antisense strands and red triangles indicate the *sfx* sites. For each panel, 4000 library members were randomly sampled and pooled from two biological replicates.

In this study we identified and tested SIs homologous to *Rci*. We established a standardised system for comparing the recombination or shuffling capabilities of these enzymes when paired with their respective recognition sites based on the ONT sequencing platform. Two such invertases, *SI-K. radicincitans* and *SI-P. punjabense*, were found to outperform *Rci* when expressed inducibly. Our findings thus expand the toolbox for shufflon-like systems in synthetic biology. *SI-K. radicincitans* is the most active shufflon invertase tested so far. It is able to randomise a 5 module shufflon during a 5-hour incubation and it recognizes an *sfx* site that differs from that of *Rci* at two nucleotide positions in both the core and right arm sequences. However, as our experiments on orthogonality demonstrate, *sfx* site variants can have a large effect on shuffling rates. Homolog *SI-E. E76* was found to exhibit a comparable shuffling activity to *Rci* with two different non-cognate *sfx* sites belonging to *SI-E. cloacae* and *SI-B. ubonensis*. Thus, while for each new SI we have based our choice of *sfx* site on the most common site found within the respective shufflon, this does not rule out that other variant recognition sites could lead to higher shuffling rates. Also, while we have tested all SI’s at 37°C, some homologs might have different optimal working conditions related to their host organism. For example, *P. parmentieri* does not survive temperatures above 33°C^19^. *H. paralvei* grows optimally at 30°C and pH 6.0^20^. To further increase the performance of the shuffling systems we describe, the parameter space around *sfx* site variants and conditions requires further mapping. The enzyme kinetics and the exact inversion mechanism of even the original *Rci* have not been fully elucidated. It is also still unclear how exactly the left arm of an *sfx* site impacts the inversion frequency. The standardised reporter assay, SIs and *sfx* variants we describe here will aid this effort. Further performance improvements may be required when one considers the shuffling of synthetic shufflons with dozens or even hundreds of modules where the ratio of *sfx* binding sites to active SI molecules may be shifted unfavourably. The performance of the enzymes we describe here should also be evaluated in eukaryotes, given that *Rci* mutants show a somewhat reduced performance in bacteria in our assays. In summary, *SI-K. radicincitans, SI-P. punjabense, SI-E. E76*, and *SI-B. ubonensis* can perform *in vivo* shuffling in *E. coli* under standard growth conditions and without the need of any co-factors, and are performing significantly better than *Rci*. Therefore, this study expands the invertase toolkit for lossless recombination employed to generate sequence diversity. It also paves the way for further discovery and testing of other such invertases which could eventually lead to establishing a set of fully orthogonal SI systems.

## Materials & Methods

### Plasmid construction

The sequence of all expression and reporter plasmids is provided in Supplementary File 2 and all primers used for cloning are provided in Supplementary Table 1. Briefly, *Rci* and its homologs were obtained by gene synthesis (ATUM, Newark, CA) in the low-copy Electra MOTHER™ pM279 backbone including an AatII restriction site and pLacIQ promoter upstream, and an XhoI restriction site and T1 terminator downstream. For inducible expression, the plasmid backbone was first obtained via a restriction digest of one of the constitutive expression plasmids using XhoI and AatII restriction enzymes which removed the pLacIQ promoter and the SI. The L-arabinose operon was then PCR-amplified using the ‘AraC-pBAD FWD’ and ‘AraC-pBAD REV’ primers. *Rci*, and homologs *SI-K. radicincitans, SI-P. parmentieri* and *SI-P. punjabense* – which were delivered by ATUM as PCR products, were PCR-amplified using the forward primer ‘Homolog FWD universal’ and the reverse primers ‘Inducible Rci EXP REV’, ‘K. rad. EXP REV’, and ‘P. parm. & P. punj. EXP REV’ respectively. Tripart Gibson assembly was then carried out using the backbone, the L-arabinose operon, and the SI fragments. For the reporter plasmids, five dummy sequences were PCR-amplified from the CDS of the transcription factor p65 activation domain, the Epstein Barr virus R transactivator activation domain, and eGFP in an expression vector in the Windbichler lab. Each variant *sfx* site was introduced via overhangs of these primers. Modules were PCR-amplified with the following primer pairs: SI-specific FWD1 and SI-specific REV1 for module 1; ‘Universal FWD2’ and SI-specific REV2 for module 2; ‘Universal FWD3’ and SI-specific REV3 for module 3; ‘Universal FWD4’ and SI-specific REV4 for module 4; ‘Universal FWD5’ and SI-specific REV5 for module 5 (Supplementary Table 1). Each set of five modules was then Gibson-assembled into a cloning vector encoding an ampicillin resistance marker, ColE1 origin of replication and ROP gene.

### Shuffling assay

One Shot™ Stbl3™ Chemically Competent *E. coli* (Invitrogen) cells were co-transformed with 3 ng DNA of both expression and reporter plasmids for an average molar ratio of 1:1.7, following the dedicated heat-shock protocol. After cell recovery, 6.8 mL of LB containing 100 μg/mL ampicillin and 34 μg/mL chloramphenicol were inoculated with the total volume of transformed cells, and incubated for 24 hours in a shaking incubator, at 37°C and 225 rpm. For inducible SIs, overnight cultures were used to seed a fresh culture at an OD600 of 0.1 which was incubated as before until an OD600 of 0.5 units. L-arabinose was then added to each culture for a final concentration of 0.2% followed by another 5-hour incubation. DNA was extracted using the QIAprep Spin Miniprep Kit (QIAGEN) with the following modifications for the 24-hour cultures. P1, P2, and N3 reagent volumes were doubled, and the cell lysis was thus carried out in 2 mL Eppendorf tubes. After the 10 min spin, the supernatant was loaded in roughly even parts on two columns. Finally, the DNA was eluted in 50 μL 53°C DNase/RNase-free Distilled water (ddH_2_O) from each column, and the two eluates were subsequently pooled to yield roughly 100 μL of DNA per sample.

### Library preparation and nanopore sequencing

Plasmid DNA was digested using NdeI (NEB) and following gel electrophoresis the 4.5kb band corresponding to the shufflon was extracted for each sample and gel-purified using Monarch® DNA Gel Extraction Kit (NEB). 200 fmol of linearised DNA per condition were barcoded using the Native Barcoding Kit 24 (Q20+) (SQK-NBD112.24, Oxford Nanopore Technologies). Sequencing was performed using a Flongle Flow Cell R9.4.1 (FLO-FLG001), and the MinION Mk1B device. 20 fmol of the pooled library was sequenced until 150,000-200,000 reads were obtained. Basecalling was performed using Guppy basecaller^21^ via the Imperial College Research Computing Service^22^, with the super accurate model (dna_r9.4.1_450bps_sup.cfg), deconvoluting reads according to barcodes (barcode_kits SQK-NBD112-24), and with trim_barcodes and trim_adapters flags on.

### Bioinformatic analysis

The analysis pipeline is found here https://gitfront.io/r/jkatalinic/xf8oSaYQpkmC/Rci-Variants/. Briefly, for each reporter the set of 384 references was generated including ∼2 kb anchor sequences flanking the shufflon. Sequencing reads were size-filtered using Seqkit^23^ for DNA of the approximate size of the shufflon and anchor sequences (4.5 kb) and mapped against the reference set using Minimap2^24^. Samtools^25^ was used to generate one compressed .bam file per reference which is filtered for alignment quality (AS score of >7500)^24^. From each filtered .bam file, a text file that reflects the order of modules is generated for each read. These files are then compiled and analysed in R.

### Protein modelling

Models for the following SIs were sourced from AlphaFold Protein Structure Database: *Rci* (AF-P16470-F1), *SI-E. E76* (AF-A0A5J6W1B6-F1), *SI-K. radicincitans* (AF-A0A1V0LM40-F1), *SI-M. glucosetrophus* (AF-C6XEL1-F1), *SI-P. citronellolis* (AF-A0A1A9KP24-F1), *SI-P. parvum* (AF-A0A6P1S1×8-F1). Models for the other 9 homologs were generated using the protein sequences from Supplementary File 1 via AlphaFold3 server^26^. Figures were generated in ChimeraX^27^. Cre was taken from PDB file 1KBU.

### Statistics

All tests were conducted in R. For comparing the shuffling rates of *Rci*, its homologs, and mutants, Welch’s ANOVA was performed for a one-way analysis of means of unequal variance^28^ to determine if the shuffling rate means are different. Unequal variance was confirmed using Bartlett test of homogeneity of variances^29^. Dunnett’s C test was performed for pairwise comparisons to a control, in this case *Rci*, in order to determine which SIs produced a mean shuffling rate of statistically insignificant difference to *Rci* for constitutive, and significant difference for inducible homologs. For assessing the correlation between module inversion frequency, module length and SI, a regression analysis was carried out in R using a linear model. For the test of orthogonality, we used a two-sample t-test with Welch’s correction since unequal variance was indicated by the difference in standard deviations between groups.

## Author Contributions

**JK**: Conceptualization, Investigation, Software, Formal Analysis, Visualization, Writing - Original Draft Preparation, Supervision. **MR**: Investigation, Software. **AA**: Methodology, Investigation. **JHM**: Data Curation. **MK**: Conceptualization **NW**: Funding Acquisition, Conceptualization, Supervision, Writing - Review & Editing.

## Supporting information

Supporting information: protein sequences of Rci homologs (.fasta), expression plasmid and reporter plasmid maps as GenBank files (.gb).

## Acknowledgments

The authors thank Alex Ivanov. The coding sequences of *Rci* mutants were kindly provided by Yi Wu from School of Chemical Engineering and Technology, Tianjin University, China.

---

**Supplementary Table 1:**
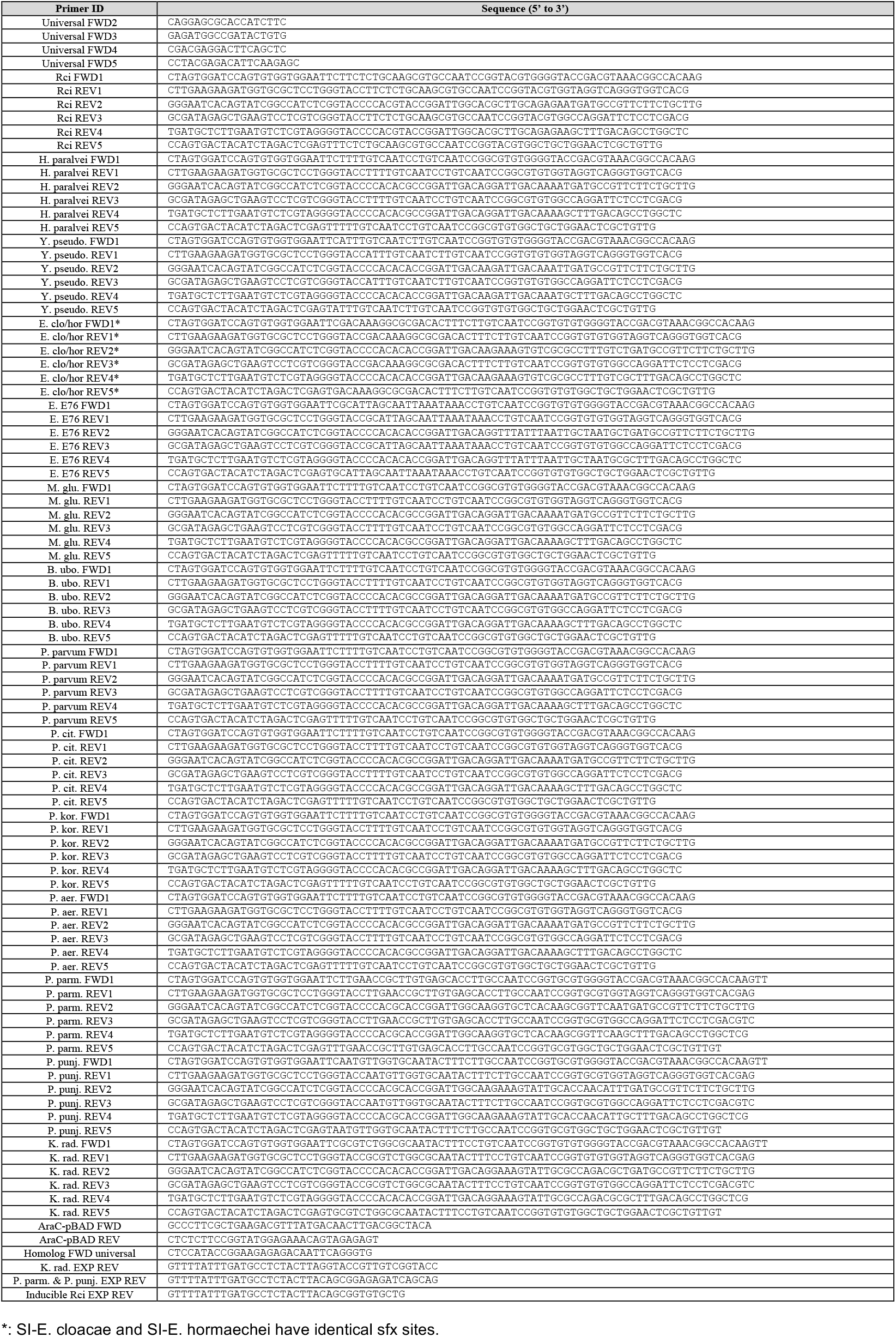
Reporter and inducible expression vector assembly primers.

**Supplementary Figure 1.**
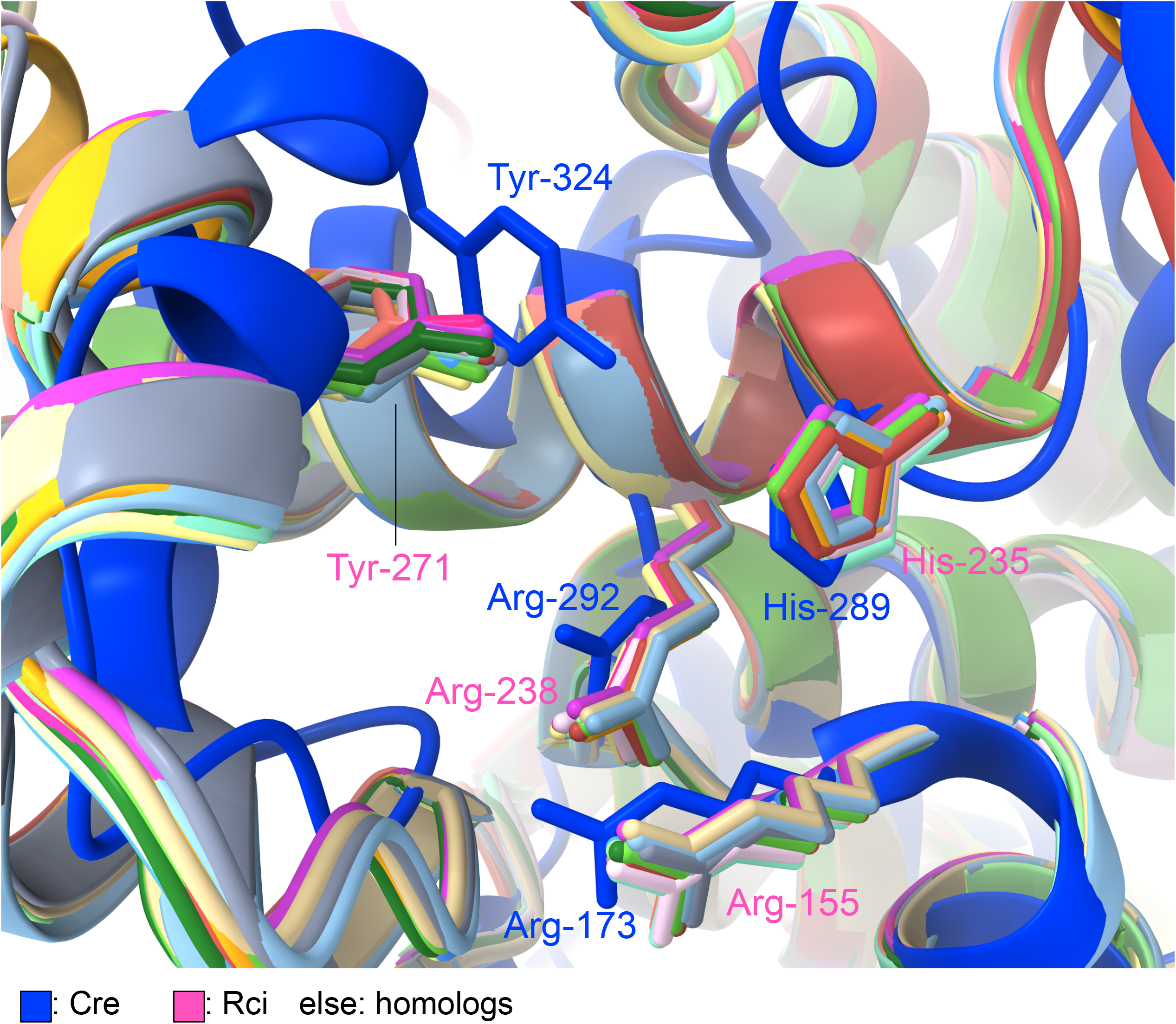
Conservation of catalytic residues in Rci homologs. Catalytic tetrad Arg-His-Arg-Tyr displayed in stick style in an overlay of protein models of *Rci, Rci* homologs, and the protein structure of *Cre*.

**Supplementary Figure 2.**
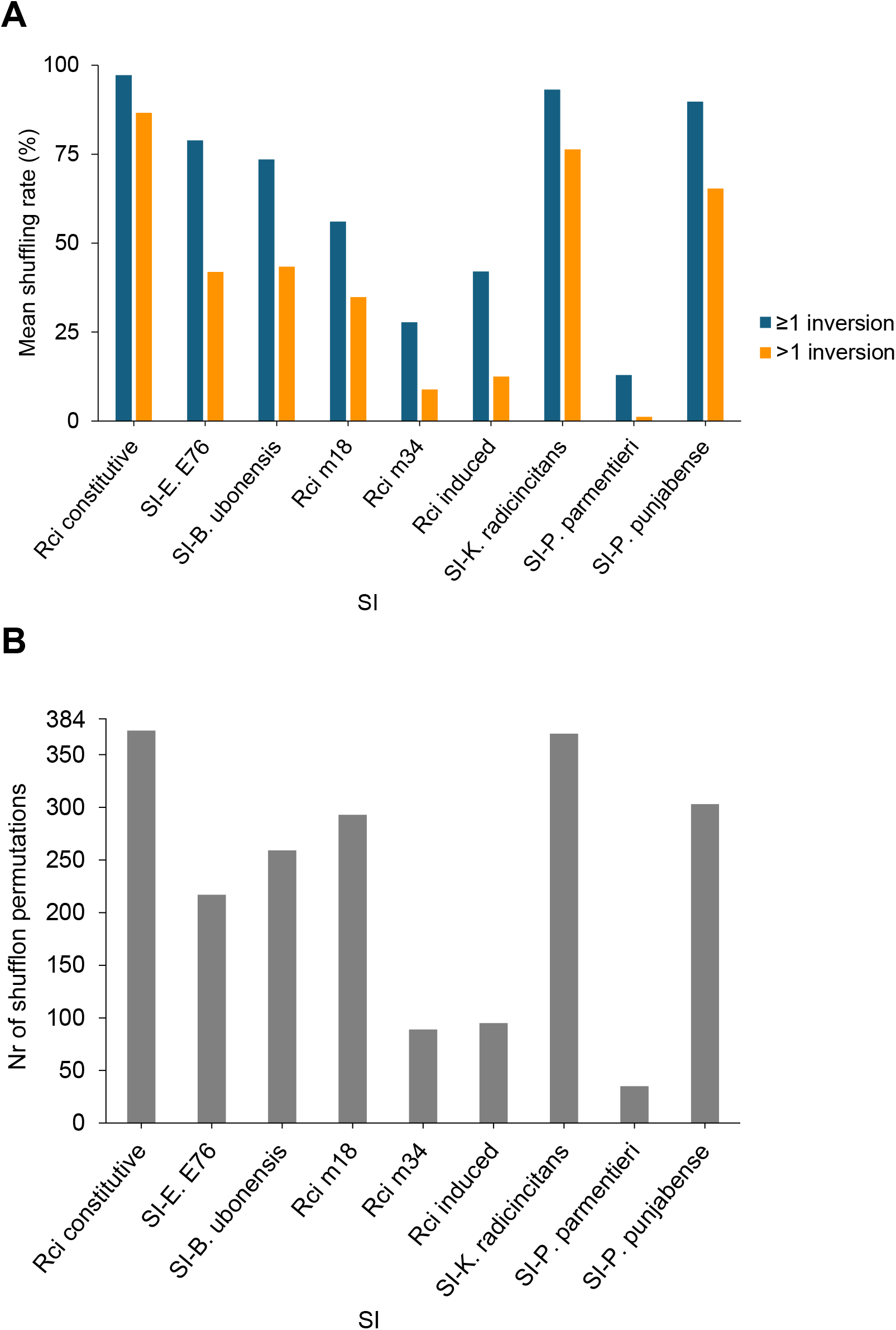
Analysis of shuffled libraries. **A**. For each SI, the mean shuffling rate (orange) which measures the proportion of analysed library members which contain shufflons that are not in the original (unshuffled) configuration as well as the multiple-shuffling rate (blue) which measures the proportion library members that are the product of more than 1 inversion are shown for the pooled data of both biological replicates. **B**. Permutations were quantified from 4000 randomly sampled library members for each SI from the pooled data of both biological replicates. There are a maximum of 384 possible permutations for a five-module shufflon.

**Supplementary Figure 3.**
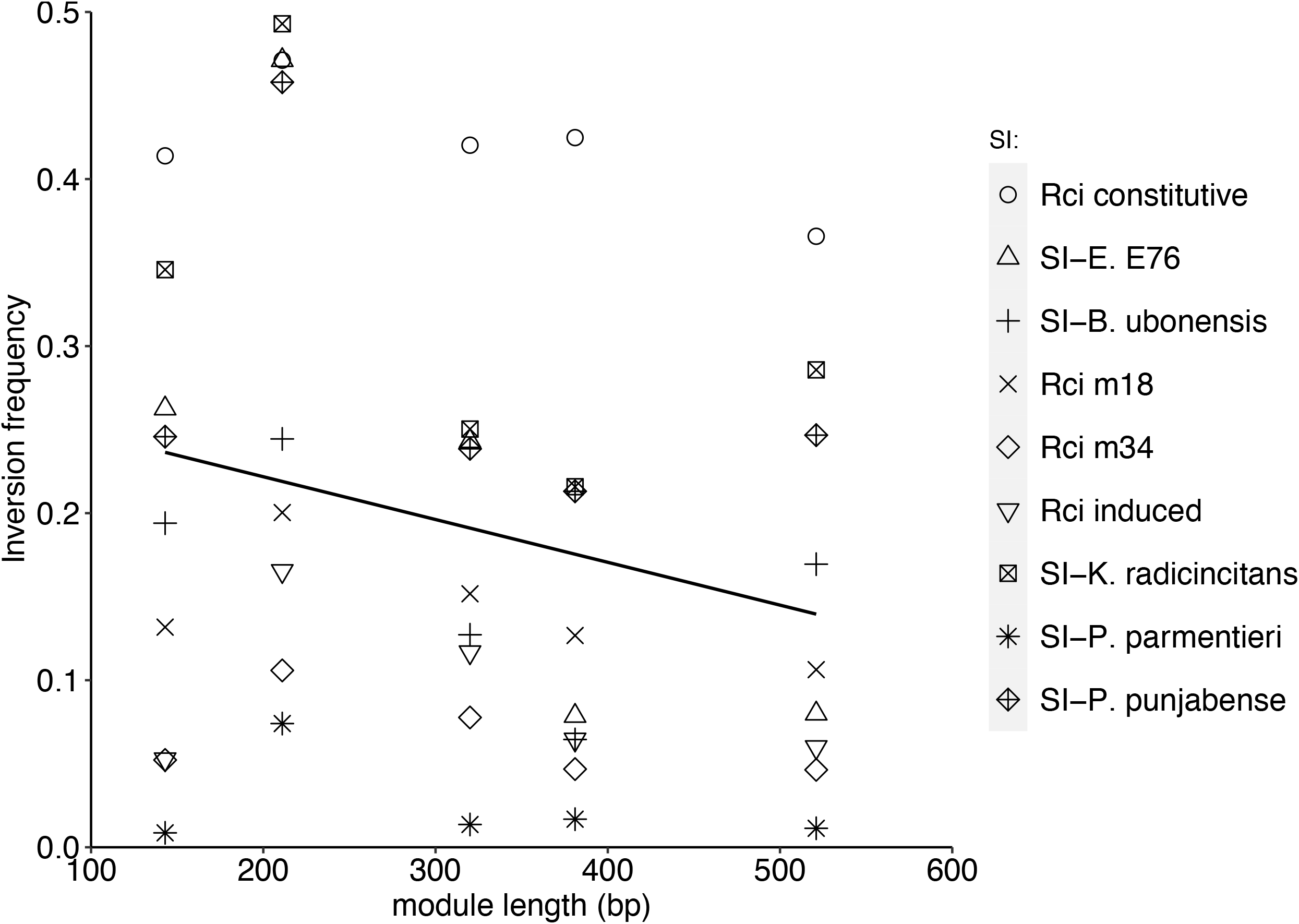
Analysis of single module inversions. For each SI, those library members from both pooled biological replicates that contained the reference module configuration (in either inverted or non-inverted form) were extracted and then randomly downsampled to the smallest library of 2201 members. A multiple regression analysis indicated that module length (p<0.00361) and SI (p<1.484×10^−9^) were regression coefficients significantly explaining the module inversion frequencies.

